# Identification of traits and functional connectivity-based neuropsychotypes of chronic pain

**DOI:** 10.1101/421438

**Authors:** Etienne Vachon-Presseau, Sara E. Berger, Taha B. Abdullah, James W. Griffith, Thomas J. Schnitzer, A. Vania Apkarian

**Affiliations:** Department of Physiology; Department of Medical Social Sciences; Departments of Internal Medicine and Rheumatology; Department of Physical Medicine and Rehabilitation; Department of Anesthesia, Northwestern University Feinberg School of Medicine, 710 N Lake Shore Drive, Room 1020, Chicago, IL 60611, USA; Healthcare and Life Sciences Department, IBM Watson Research Center, 1101 Kitchawan Rd, Yorktown Heights, NY 10598, USA

**Keywords:** Personality, psychology, fMRI, connectomics, catastrophizing, socioeconomics

## Abstract

Psychological and personality factors, socioeconomic status, and brain properties all contribute to chronic pain but have essentially been studied independently. Here, we administered a broad battery of questionnaires to patients with chronic back pain (CBP). Clustering and network analyses revealed four orthogonal dimensions accounting for 60% of the variance, and defining *chronic pain traits*. Two of these traits – *Pain-trait* and *Emote-trait* - were related to back pain characteristics and could be predicted from distinct distributed functional networks in a cross-validation procedure, identifying *neurotraits*. These neurotraits were relatively stable in time and segregated CBP patients into subtypes showing distinct traits, pain affect, pain qualities, and socioeconomic status (*neuropsychotypes*). The results unravel the trait space of chronic pain leading to reliable categorization of patients into distinct types. The approach provides metrics aiming at unifying the psychology and the neurophysiology of chronic pain across diverse clinical conditions, and promotes prognostics and individualized therapeutics.

---

Unraveling the mechanisms of chronic pain remains a major scientific challenge. There is now strong and convincing evidence that specific brain properties contribute to the risk of developing chronic pain, and that the transition to chronic pain involves brain adaptations that, to a large part, construct and mold the state of chronic pain (Baliki and Apkarian, 2015; Vachon-Presseau et al., 2016). Additionally, current theories suggest that pain characteristics, pain-related disability, and responses to treatment are all partially determined by psychological factors and/or personality properties (Turk and Okifuji, 2002), as well as parameters related to socioeconomic status (SES) (Green and Hart-Johnson, 2012). Although the biopsychosocial (BPS) perspective is actively applied in the everyday clinical management of chronic pain (Kamper et al., 2014), component properties that comprise the concept – namely biological processes (brain and body), psychological factors, and the impact of social factors on these components – have not been jointly studied. Thus, the relative influence of these factors on each other, as well as their independent contribution to the state of chronic pain, remains unknown. In its current form, the BPS model is built on fragmented evidence taken from different disciplines, and – remarkably – lacks integration between psychosocial components and underlying biology. Here, we unravel and interrelate components of BPS by combining psychological, personality, and SES factors with measures of resting state functional connectivity to begin to define a unified perspective of chronic pain.

There is a large body of literature demonstrating that psychological and personality factors are important contributors to chronic pain (e.g., the psychological components of BPS). Pain catastrophizing (Sullivan et al., 2001) and fear of pain (Goubert et al., 2004) represent strong predictors of chronic pain, whereas pain acceptance is associated with resilience (McCracken, 1998). Moreover, neuroticism – one factor in the five-factor model of personality (Goldberg, 1992) – has been shown to moderate the association between catastrophic thinking and pain vigilance (Goubert et al., 2004) and has been associated with substance abuse, anxiety, and major depressive disorder (Griffith et al., 2010; Kotov et al., 2010). In contrast, optimism is usually associated with lower catastrophic thinking, higher pain acceptance, and better coping strategies (Goodin and Bulls, 2013). Other psychological factors such as mindfulness or body awareness have also been linked with improvements in pain, mood disturbance, and disability (Kabat-Zinn et al., 1985; McCracken et al., 2007), as well as with better coping strategies for managing chronic pain (Mehling et al., 2011). It is thus unsurprising that psychological factors are usually considered better predictors of chronic pain disability than the actual injury itself (Crombez et al., 1999). Despite their pivotal role in chronic pain, a comprehensive multidimensional characterization of psychological and personality factors in chronic pain remains unexplored. The first aim of the current study was therefore to determine how the psychological factors believed to impact persisting pain interacted with general personality factors to give rise to *chronic pain traits*.

The neurobiological underpinnings of chronic pain psychology/personality also remains relatively unknown, as few studies have examined the association between such factors (usually focusing on pain catastrophizing) and brain functional properties. For instance, pain rumination has been associated with stronger functional connectivity between the medial prefrontal cortex and regions within the default mode network (DMN) in temporomandibular disorder (Kucyi et al., 2014). In migraine, seed-based analyses show that pain catastrophizing is associated with functional coupling between several brain regions (posterior cingulate cortex with the dorsolateral prefrontal cortex, and the insula with the hippocampus) (Hubbard et al., 2014). In fibromyalgia, pain catastrophizing has been associated with increased dorsolateral prefrontal cortex activity in anticipation of pain (Loggia et al., 2015). However, these studies tested limited psychological parameters in association with brain properties from only a few regions-of-interest. Here, our secondary aim was to decode traits of chronic pain using resting state functional connectivity magnetic resonance imaging (rsfMRI), with the goal of determining *if* and *how* chronic pain traits are represented in patterns of brain activity, bridging the psychology of pain with a biological framework, which we define as *neurotraits*. Thus, our primary hypothesis was that chronic pain traits can be predicted from distributed neural representations. Additionally, we posited that mapping the neural representation signatures back onto the multidimensional trait and pain characteristics spaces could, in turn, establish the existence of subtypes of chronic pain patients. This latter step would lead to the finding and characterizing a neurobiologically-informed psychological/personality typology, which we label *neuropsychotypes* of chronic pain.

Our third aim was to determine the influence of social components on traits, neurotraits, and neuropsychotypes of chronic pain by examining the impact of socioeconomical status (SES) on these dimensions. Many studies show that lower SES is associated with higher prevalence of musculoskeletal pain, neuropathic pain, ulcers, and sciatica across Europe and North America (Poleshuck and Green, 2008). Consistently, the British Birth Cohort study revealed that lower SES during adulthood is associated with shoulder, forearm, low back, knee, and chronic widespread pain (Macfarlane et al., 2009). Consistent with the BPS model, SES and occupational hazards are dominant risk factors for the onset of back pain and pain disability, independent of injury occurrence or type (Dorner et al., 2011; Katz, 2006). We therefore hypothesized that SES measured with self-reported income, years of education, gender, and ethnicity will influence the traits, underlying neurobiology, and impact on neuropsychotypes of chronic pain.

Our results indicate four dimensions comprising *chronic pain traits*, only two of which were associated with back pain characteristics of CBP. We therefore studied brain functional connectivity networks for these two dimensions and demonstrate that they can be predicted from three distributed functional networks (*neurotraits*). These networks were relatively stable in repeat brain scans and mediated changes seen in pain, anxiety, and negative affect. The neurotraits were then used to derive five clinically-meaningful neuropsychotypes of CBP patients with specific pain characteristics. Confirming our starting hypothesis, SES influenced specific chronic pain traits and their related functional networks, and were different across identified neuropsychotypes.

## Results

This study is based on data collected in the setting of a Randomized Control Trial (RCT) designed to study placebo pill response in chronic pain patients (Vachon-Presseau et al., 2018). Patients with chronic back pain (CBP) enrolled in the RCT visited the lab on six occasions with four visits including brain-imaging sessions (**fig. 1a**). A total of 108 CBP patients were included. Group 1 included 62 CBP that completed all 6 visits of the RCT and were used to determine the components of personality, derive underlying patterns of brain connectivity, and cluster the patients into neuropsychotypes. Group 2 included 46 CBP patients that completed all questionnaires at V1 but did not complete the brain imaging sessions for various reasons, including claustrophobia, meeting an exclusion criterion at a later point in the trial, or dropping out of the study. The demographics for these patients can be found in **supplementary table 1**.

**Fig. 1.**
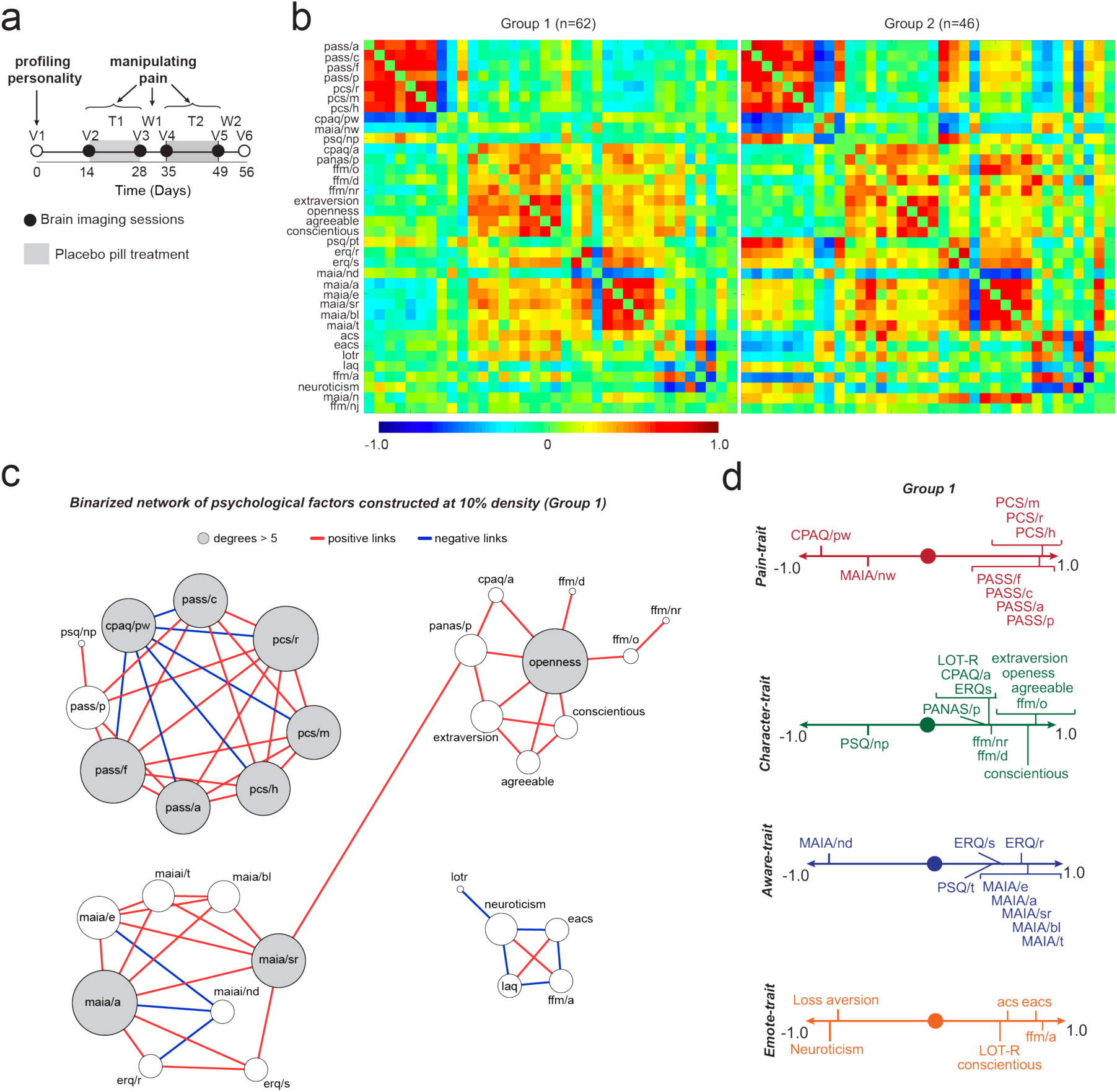
Psychological and personality factors identify four chronic pain traits. **a.** The study design consisted of six visits to the lab with four of these including brain imaging sessions. Questionnaires were administered at V1 and pain was manipulated with placebo treatment (T1) followed by a washout period (W1), repeated twice (T2, W2). 62 patients completed all 6 visits (Group 1) and 46 patients completed the questionnaires at V1 but did not finish the rest of the study (Group 2). **b.** Each covariance matrix (Group 1 and Group 2; ordered based on PCA results for Group 1) showed associations between the questionnaire outcomes used to probe CBP patients’ profiles (r-values shown in bar from blue to red). **c.** A network constructed from the Group 1 covariance matrix displays the strongest correlations (top 10%) and topological properties indicated the presence of 4 communities. **d.** PCA performed on all 36 questionnaire outcomes (Group 1) identified 4 orthogonal components. Given the observed loadings and network clusterings, we label these components as: *Pain-trait*; *Character-trait*; *Aware-trait*; and *Emote-trait*. Only components loading > 0.4 or < −0.4 are depicted. For each figure, a description of the analyses and p-values are reported in Supplementary Table 4.

### Psychology/personality traits of chronic pain

We first sought to define the psychology/personality factors of chronic pain based on responses to a broad battery of questionnaires in two CBP cohorts (Group 1 and Group 2; **supplementary table 1**). We applied clustering and network analysis methods to the covariance matrix constructed from responses to questionnaires that surveyed concepts related to chronic pain, constructs of personality, and emotional, attentional, and motivational properties, all of which are thought to influence traits of chronic pain. We profiled the CBP patients using the following measures: pain anxiety (PASS), pain catastrophizing (PCS), pain sensitivity (PSQ), pain acceptance (CPAQ); emotion regulation (ERQ), attentional capacities (ACS), attentional capacities in the presence of emotions (EACS), optimism (LOT-R), sensitivity and aversion to loss (LAQ), positive affect (PANAS/p), interoceptive awareness capacities (MAIA), mindfulness capacities (FFM), and personality traits (NEO-5: neuroticism, extraversion, openness, conscientiousness, and agreeableness). The associations among subscales are presented in **fig. 1b** for both groups of patients. PCA was used to order the 36 subscales in Group 1 and subsequently used to organize the questionnaires for Group 2 in **fig. 1b**. As shown in the figure, the pattern of covariance was very similar in the two groups.

We used Group 1 data to construct a network from the covariance matrix of 36 subscales. This network exhibited high-dimensional topology, as community clustering of the subscales segregated into four distinct communities (**fig. 1c)**. PCA analysis applied on the covariance matrix for Group 1 also identified four orthogonal principal components with eigenvalues > 2.0 explaining more than 5% of the variance (representing the elbow in the scree plot, **supplementary fig. 1a**). The four components together explained 63% of the variance. PCA performed for Group 2 data identified the same four components as seen for Group 1 data, explaining 57% of the variance, validating the four dominant psychological dimensions related to chronic pain. These components are dubbed “*chronic pain traits*,” which we label based on the component loadings. Component 1, *Pain-trait*, was dominated by factors emphasizing the toll of chronic pain: high positive loadings for fear of pain, pain catastrophizing, and pain anxiety, and a high negative loading for pain acceptance. Component 2, *Character-trait*, included mainly positive personality traits such as extraversion, conscientiousness, agreeableness, and openness. Component 3, *Aware-trait*, included various awareness capacities and abilities to regulate emotions. Component 4, *Emote-trait*, included diminished neuroticism, lower sensitivity to loss, high optimism, strong attentional control in the presence of emotion, and high aptitude for mindfulness. Similar results were obtained in Group 2, where the PCA identified the same four domains with factor loadings similarly, as seen in Group 1 (**supplementary fig. 1b**). Although network and PCA analyses resulted in similar results, there were also subtle differences in the components identified by each procedure, thus we use both representations in subsequent analyses. Overall, we identify four validated dimensions comprising traits of chronic pain in CBP patients. Unsurprisingly, the obtained pattern indicates that pain-related properties comprise the dominant dimension, while the other three dimensions comprise positive domains of character, awareness, and emotion, all of which would theoretically counterbalance or modulate the primarily negative pain-related dimension.

### Influence of chronic pain traits on back pain characteristics

We examined the interrelationship between the four trait dimensions of chronic pain with the intensity, qualities, and negative affective characteristics of the back pain experienced by our CBP patients. Our aim was to link specific traits to pain aspects of CBP. In both groups, we assessed back pain and pain-related negative affect using questionnaires and ratings. The pain intensity measurements included a series of numerical scales: out-of-clinic ecological momentary assessments (EMAs) of back pain intensity entered twice a day using smartphone technology, a verbal recall of the pain experienced over the last seven days (Memory), and an in-lab numeric rating scale of the current pain (NRS). Pain qualities were assessed using the McGill pain questionnaire – sensory (MPQ/s) and affective (MPQ/a) subscales – and PainDetect. Negative mood and depressive symptoms were assessed using the negative affect score from the Positive and Negative Affect Scale (PANAS/n) and the Beck Depression Index (BDI-1a). Finally, physical health was also evaluated (SF-12p). The absolute levels in these pain measurements were equivalent between the two groups with the exception of SF-12p that was higher in Group 1. Baseline levels for these measurements are presented for both groups in **supplementary table 2** and their covariance is presented in **supplementary fig. 2**.

The correlation matrix between trait dimensions and pain measurements was calculated for visit 1 and visit 2 in Group 1 (**fig. 2a**), and for a smaller set of pain measurements for visit 1 in Group 2 (**fig. 2b**). Obtained results indicate that *Pain-trait* was positively related to almost all clinical measurements capturing back pain properties at visit 1 (V1) and visit 2 (V2), while *Emote-trait* was anticorrelated primarily to the negative affective characteristics of CBP (**fig. 2a**, Group 1). Similar associations were also observed in Group 2 (**fig. 2b**). *Character-trait* and *Aware-trait* showed no reliable relationships with pain or negative affect measurements. Consensus analysis between visits and groups confirmed these results (**fig. 2c**). Thus, here we uncover two traits influenced specifically by back pain properties, in opposite directions and with distinct patterns.

**Fig. 2.**
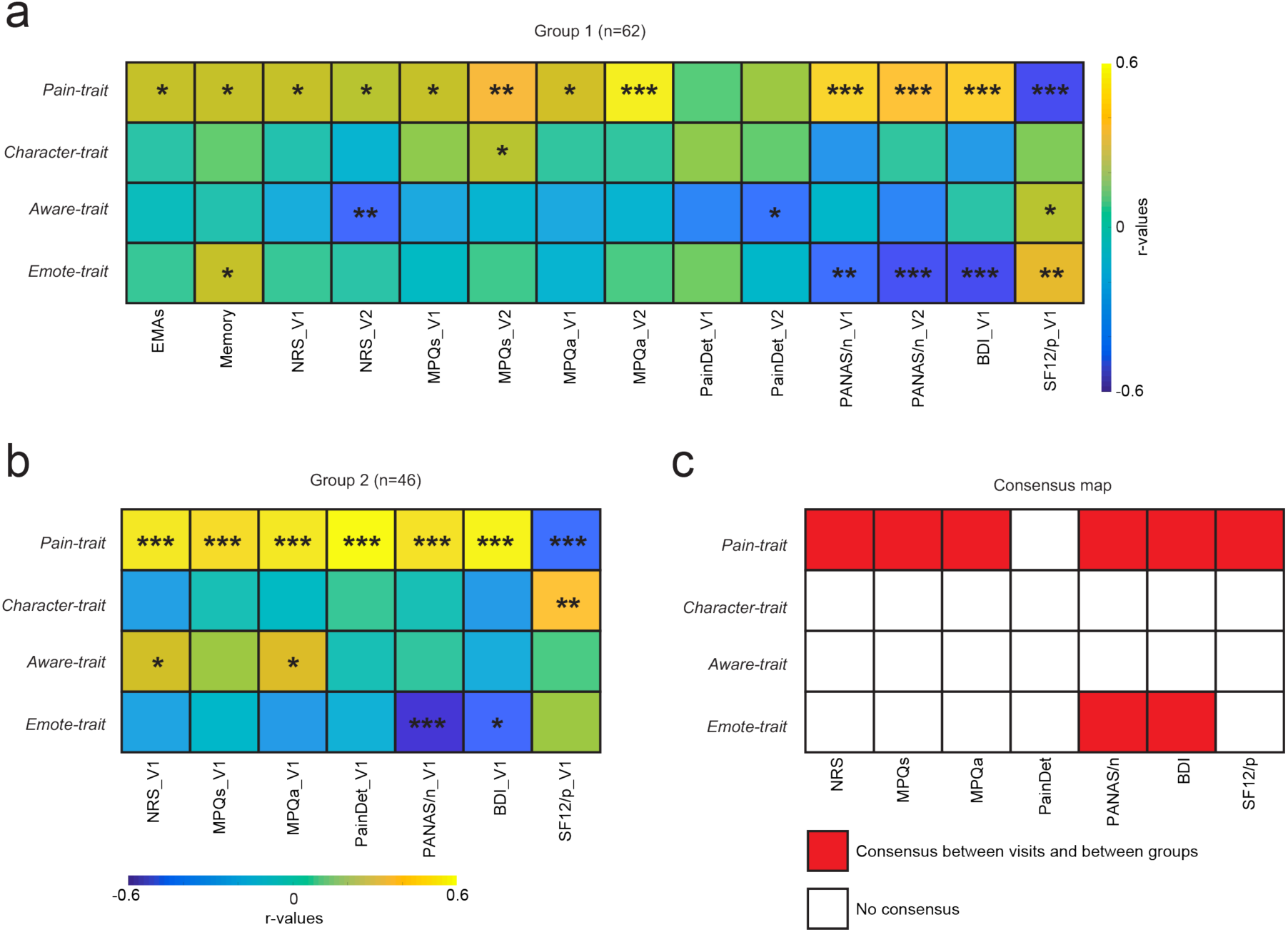
Relating trait dimensions to pain characteristics. Correlation matrices indicate the extent of influence of each trait with back pain intensity (EMAs, Memory, NRS), quality (MPQs/a, PainDetect), and negative affect (PANAS/n, BDI). **a.** In group 1, *Pain-trait* scores were related to almost all pain characteristics queried at visit 1 (V1) and visit 2 (V2), while *emote-trait* scores were negatively related to negative affect at V1 and V2. **b.** Consistent findings were observed in group 2 (data only available at V1). **c**. Consensus was obtained when the effects were replicated within participant (across V1 and V2 in Group 1) and between cohort of patients (Group 1 and Group 2). The consensus map shows that *Pain-trait* related to almost all pain assessments and *Emote-trait* related specifically to negative affect. BDI: Beck depression index (only collected at V1); EMAs: ecological momentary assessments (phone app daily pain ratings); NRS: numerical rating scale; MPQa: McGill pain questionnaire/affective; MPQs: McGill pain questionnaire/sensory; Pain Detect; PANASn: positive and negative affect schedule/negative; SF12/p: 12-item short-form survey/physical health. The consensus across visits and groups was established using uncorrected p-values ^*^ p < 0.05; ^**^ p < 0.01; ^***^ p < 0.001 (significant even after Bonferroni correction for all comparisons (56 comparisons in Group 1 and 28 comparisons in Group 2)).

Finally, we assessed if the expression of these chronic pain traits was influenced by how long individuals have been suffering from persisting pain; we observed no effect of pain duration on any traits (Pearson correlations; all p values > 0.24).

### Brain functional circuits for neurotraits Pain-trait and Emote-trait

Next, we examined if the two chronic pain traits associated with back pain properties can be decoded from functional networks using resting state fMRI in a cross-validation procedure. We used connectome-based predictive modeling of functional connectivity to determine if brain networks underlying chronic pain traits can be identified applying the Shen et al. protocol (Shen et al., 2017). First, we computed pairwise Pearson correlation coefficients between the time courses of each possible pair of 272 nodes; these generated a single subject connectivity matrix. We then used a 3-fold cross-validation procedure, where patients were divided into training (2/3 of patients) and testing (1/3 of the patients) sets. The training set was used to select features associated with each trait score, performing robust regressions between each link of the connectivity matrices with the specific trait score (p < 0.05). A leave-one-out procedure was used within the training set to reduce the number of connections to the ones correlating with the trait in each of the n-1 iterations. Once the stable connections were identified, a summary statistic was calculated from the sum of all edges (*z(r)*) positively correlating with the trait and the sum of all edges (*z(r)*) negatively correlating with the trait, separately. No other approach was tested to predict brain circuitry for chronic pain traits.

The predictive values of these correlates were validated using the left-out patients of the test set. The sum of positive and negative z*(r)* values from the same set of edges identified in the training set was calculated in the patients of the test set to predict their traits. This cross-validation procedure was repeated for the three folds, rotating and substituting the 1/3 of the patients used in the test set. **Fig 3.a** shows that three patterns of functional connectivity were successfully cross-validated using this procedure. The *Pain-trait* score was predicted from both positive (*r*^*2*^ = 0.14; *p =* 0.003) and negative edges (*r*^*2*^ = 0.10; *p =* 0.01) and the *Emote-trait* score was predicted only from positive edges (*r*^*2*^ = 0.08; *p =* 0.03). Because the features selected in the training set differed across the three folds, a single set of consensus weights was generated from averaging the links across the folds. Links persistent across the three folds were labeled ‘most important predictors’ and displayed (**fig. 3b,d,f)**.

**Fig. 3.**
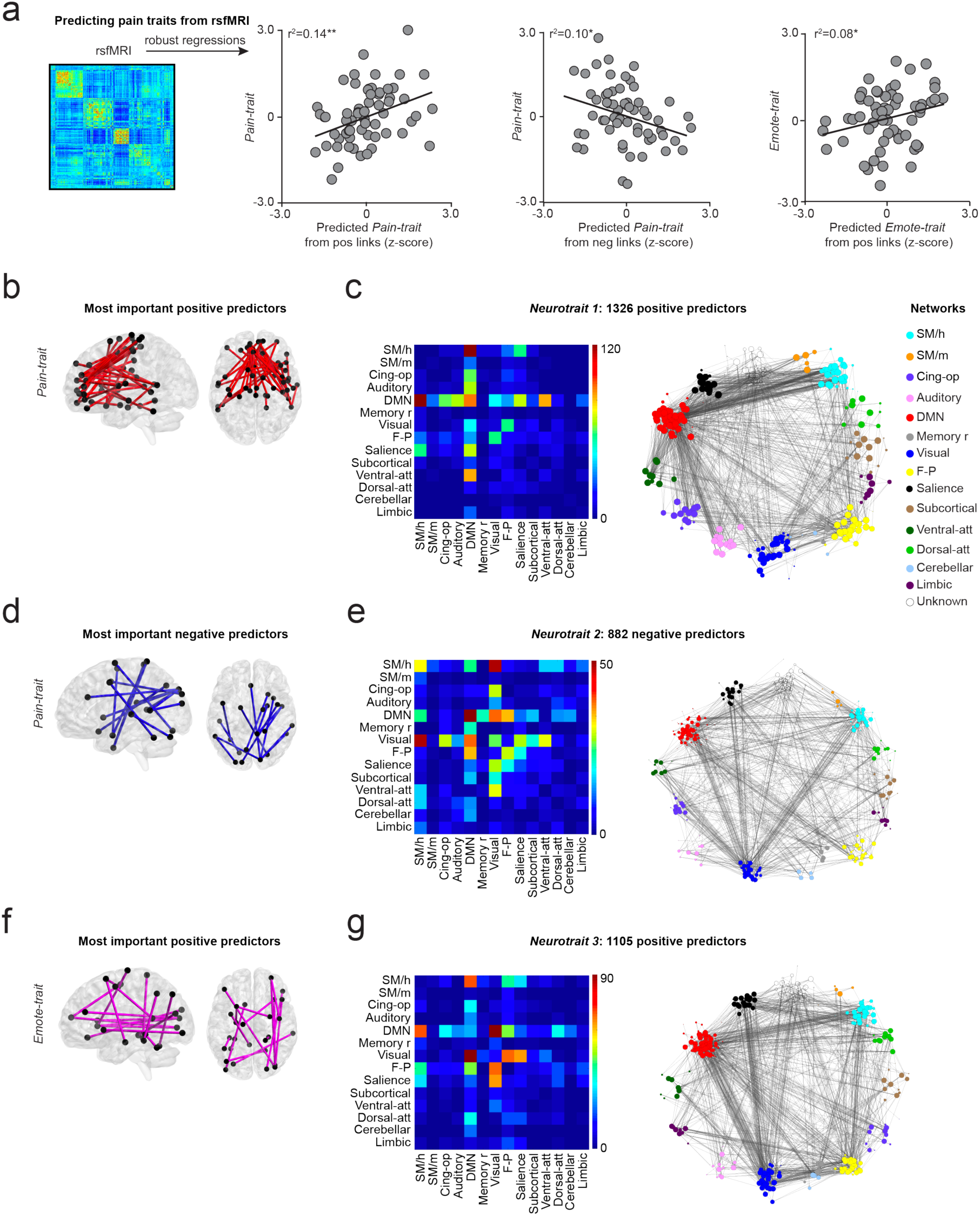
Connectome-based predictive modeling predicts *Pain-trait* and *Emote-trait*. **a.** Resting-state fMRI was used to construct functional connectivity covariance matrices, and a 3-fold cross-validation procedure was implemented to identify links related to trait scores. Scatterplots show the predicted trait score of each individual when they were in the left-out fold (predictive value in new individuals). *Pain-trait* was predicted from both positive and negative links and *Emote-trait* could be predicted only by positive but not negative links. **b-d-f.** The most important predictors (MIPs) were the common links consistently selected by the models across all 3 folds. **c, e, g.** The matrices display within- and between-community connections of all predictive links, irrespective of their weights, when the brain networks are represented by 14 communities. The full connectivity profile for each trait is also shown across all 272 nodes. ^*^ p < 0.05; ^**^ p < 0.01.

All weighted links were organized based on their connectivity within and between the 14 communities of the human brain (**fig. 3c, e, g**). The full connectivity matrices for all links related to the two traits are also shown (**fig. 3c, e, g**). *Pain-trait* scores were predicted from a total of 1,326 positive links: mainly between DMN and sensorimotor, cingulate, salience, and ventral attention communities (*Neurotrait 1*); and 882 negative links: mainly between sensorimotor, DMN, memory, visual, frontoparietal and ventral attention communities (*Neurotrait 2*) (**fig. 3d)**. The *Emote-trait* scores were predicted from a total of 1,105 positive links: between DMN, visual, sensorimotor, salience, and frontoparietal communities (*Neurotrait 3*). Thus, highly distributed networks comprise the neurotraits related to the two chronic pain traits specifically reflecting CBP properties.

### Neurotraits are relatively stable and their fluctuations mediate changes in anxiety and affect

All analyses presented thus far were performed on data collected at V1 (questionnaire data) or V2 (pre-treatment brain imaging and questionnaire data). Next, we examined the stability of the three identified neurotraits by assessing functional connectivity patterns across the four longitudinal brain imaging sessions. The sum of positive and negative z*(r)* values from the same sets of edges initially identified at V2 (**fig. 4a**) was calculated for the rsfMRI data collected at V3, V4, and V5 (**fig. 4b**). The pattern of functional connectivity calculated from V3-V5 – neurotraits derived weeks after assessing the corresponding traits – successfully predicted patients’ scores on *Pain-trait* and *Emote-trait*. Thus, we observe a conserved relationship between traits and related neurotraits even when the measures derived from outcomes collected weeks apart.

**Fig. 4.**
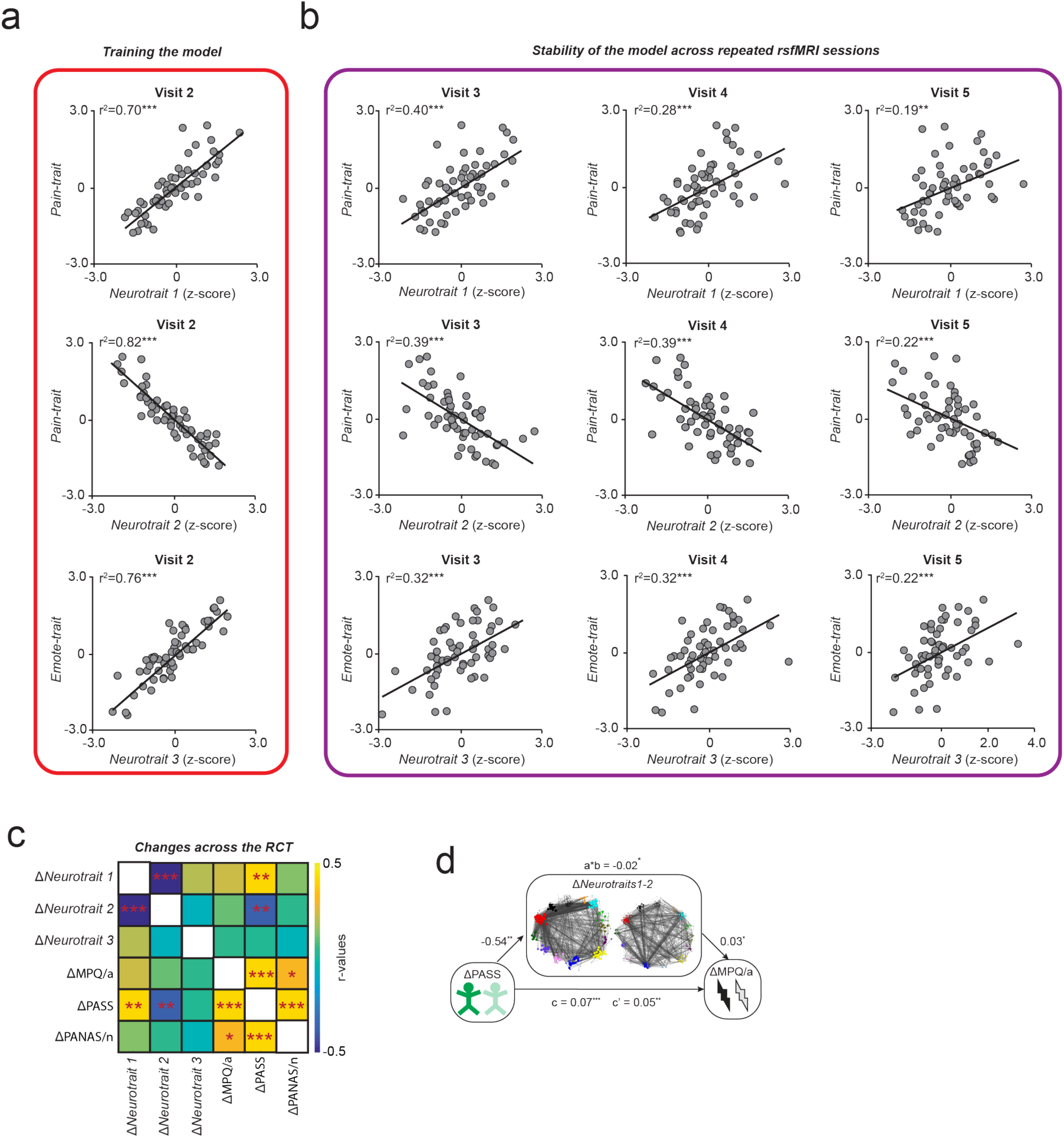
Neurotraits are relatively stable in time and their fluctuations mediate changes in pain qualities and negative affect. **a,b.** Stability assessment of neurotraits from repeat brain scans over 6 weeks. **a.** Strength of relationships between cross-validated patterns of connectivity that were trained at visit 2 with corresponding psychology/personality trait scores. **b.** Relationship between neurotraits (derived from connectivity based on subsequent brain scans) and trait scores determined by questionnaires at visit 1. The trait-neurotrait relationship is preserved. **c.** The covariance matrix shows that changes in *Neurotrait 1* (δ*Neurotrait* 1) and *Neurotrait 2* (δ*Neurotrait* 2) were associated with pain anxiety (δPASS, but not δMPQ/a, or δPANAS/n *per se*); δPASS was in turn associated with δMPQ/a and δPANAS/n. **d.** Mediation analyses showed that δ*Neurotraits 1-2* partly mediated the relationship between δPASS and δMPQ/a. ^*^ p < 0.05; ^**^ p < 0.01; ^***^ p < 0.001.

In this RCT, patients from Group 1 received either sugar pills (n=43) or no treatment at all (n=19). Pain measurements and one questionnaire measuring pain anxiety (PASS) were collected on each visit. PASS – a questionnaire strongly loading with *Pain-trait –* showed a striking diminution by about 50% between baseline V1 and post treatments V6 (δPASS, **supplementary fig. 3**). Interestingly, the δPASS correlated with decrease in pain qualities (δMPQ/s, δMPQ/a, PainDetect) and with negative affect (δPANAS/n) but not with changes in pain intensity (δEMAs, δMemory, δNRS), depression score (δBDI) or physical health (δSF12/p).

We therefore more closely examined the associations between pain anxiety (PASS), affective qualities of back pain (MPQ/a), and general negative affect (PANAS/n), which all changed over time, independent of treatment type received or presence of placebo response (**supplementary fig. 3;** see (Vachon-Presseau et al., 2018) regarding the unspecific improvement of pain qualities). We observed that changes in *neurotraits 1* and *2* between the first and the last fMRI sessions (V2 and V5; *δNeurotrait 1* and *δneurotrait 2*) were both related to δPASS, but not with δMPQ/a or δPANAS/n *per se*, suggesting that its expression pattern tracked *Pain-trait* related-outcomes rather than pain qualities or affect. These results are summarized in the covariance matrix presented in **fig. 4c.**

We then formally tested if the joint contributions of *δNeurotrait 1-2* had a causal effect on the association between δPASS and δMPQ/a, or δPANAS/n, using a series of mediation analyses. A first mediation showed that the strong relationship between δPASS with δMPQ/a at V6 (r = 0.42; p < 0.001) was partly mediated by the δ*Neurotraits 1-2* functional connectivity (**fig. 4d**). Using a three-path mediation analysis, we further showed that the changes in chronic pain were sequentially mediated by the effect of δ*Neurotraits 1-2* on δPASS. In a first model, δ*Neurotraits 1-2* mediated the relationship between MPQ/a V1 and δPASS, which in turn partially mediated the relationship between MPQ/a at V1 and MPQ/a at V6 (**supplementary fig. 4a**). In the second model, δ*Neurotrait 1-2* pattern of connectivity mediated the relationship between PANAS/n V1 and δPASS, which in turn partially mediated the relationship between PANAS/n at V1 and PANAS/n at V6 (**supplementary fig. 4b**). Importantly, reversing the order of the mediators yielded no significant results, indicating the importance of the sequence where changes in neurotraits preceded changes in δPASS. Although the effect sizes of these mediations were relatively small, they suggest that the derived functional networks may actually contribute to the affective properties of back pain.

### Defining neuropsychotypes

Here we tested if the mere expression of the neurotraits was sufficient to infer the clinical characteristics of the patients. In effect, we segregated the CBP group into subtypes based on the brain neurotrait subgroupings, thus deriving neurotrait-based types, or neuropsychotypes. Hierarchal clustering was performed on the normalized level of expression of each neurotrait-based set of brain features to derive neuropsychotypes sharing similar patterns of brain functional connectivity (**fig. 5a**). One of the challenge of this approach is to set the number of clusters to properly assess neuropsychotypes. The clustering criterion was based on a previous study showing that four types of individuals defined from personality traits characterized the general population (Gerlach, 2018). Thus, we looked for a solution around four clusters (silhouette coefficients presented for each solution in **supplementary fig. 5**). Here, the five-cluster solution was retained because it provided the finest solution segregating patients into homogeneous groups expressing distinct patterns of connectivity (**fig. 5a-b**); fairly segregated on 3D graphs representing the four trait dimensions (**fig. 5c**); and, provided clinically relevant information (**fig. 5c-h**).

**Fig. 5.**
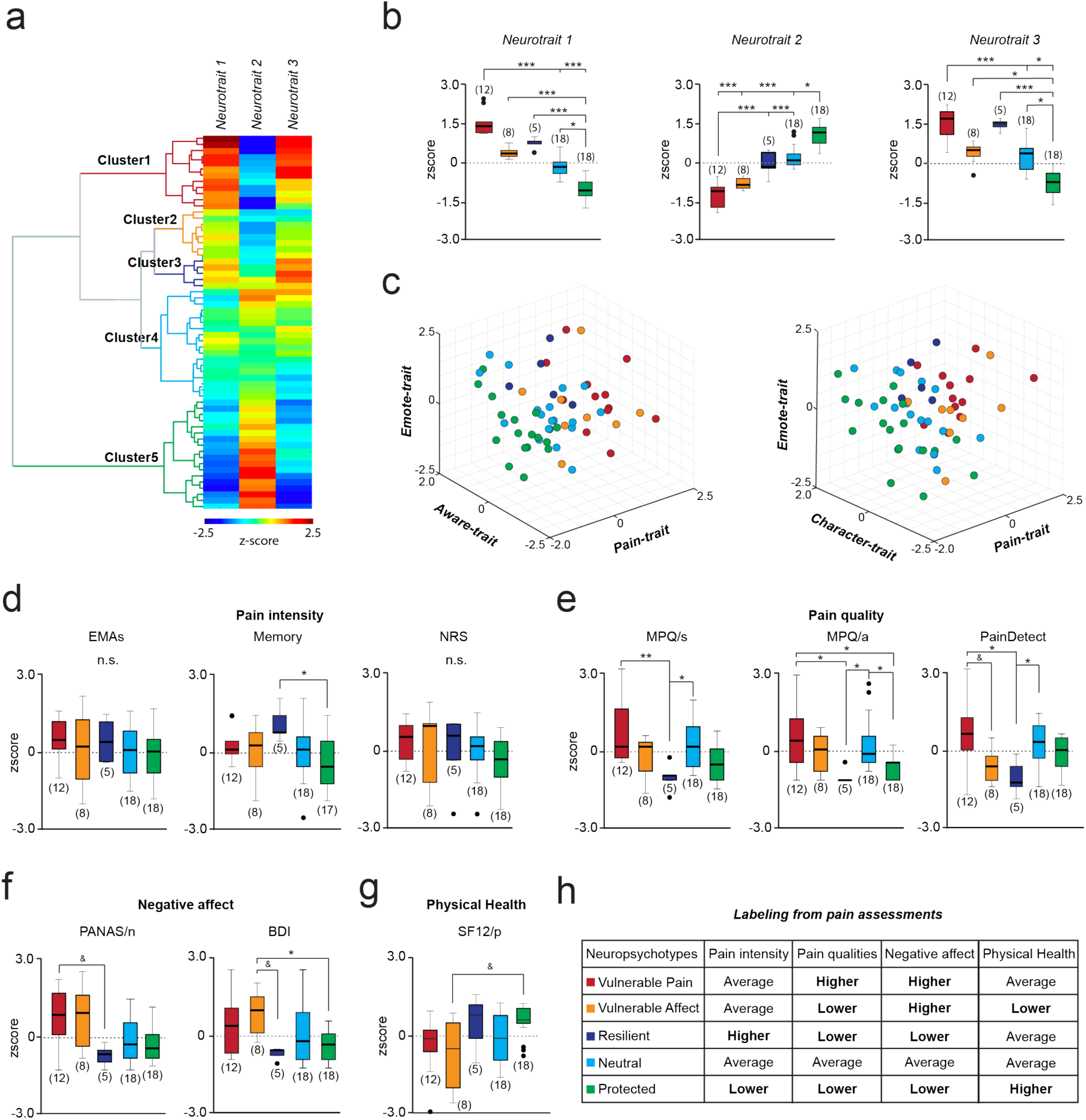
Patterns of brain connectivity, neurotraits, identified five homogenous neuropsychotypes with distinct clinical features. **a**. Hierarchal clustering showed that patients could be grouped into five different types based on the normalized expression of each neurotrait connectivity pattern, resulting in 5 neuropsychotypes. **b**. Bar graphs show relative properties of the 5 neuropsychotypes on the three neurotraits (different colored bars). **c.** Pattern of segregation for the 5 neuropsychotypes on three-dimensional rendition of the 4 trait dimensions **d-g.** The clinical profile of the 5 neuropsychotypes were assessed based on nine pain- and negative affect-related outcome measures. **d.** There were no group differences for pain intensity measured by EMAs and the numeric rating scale (NRS), but the *neuropsychotype 3* had high pain memories (verbal recall of pain experienced over the last week). Neuropsychotypes differed on: **e.** pain qualities measured with the MPQa, MPQs, and PainDetect; on **f.** negative affect measured with PANAS/n and BDI; as well as on **g.** physical health (SF12/p). **h.** In addition to traits, each neuropsychotype was also labeled based on correspondence across the nine clinical back pain outcomes.

### Characteristics of the five neuropsychotypes

We examined the characteristics of the five chronic pain neuropsychotypes using the nine back pain related measurements (EMAs, memory, NRS, MPQ/s/a, PainDetect, PANAS/n, BDI, SF12). Our results show that seven out of the nine measurements differed between the neuropsychotypes. Measures of pain intensity showed the smallest effects, where only a verbal recall of the pain experienced over the last week (memory) was different between the neuropsychotypes (**fig. 6a**). Pain qualities, negative affect, and physical health were, however, consistently different across the 5 neuropsychotypes (**fig. 6b-d**). Given these clinical differences, and also taking into account differences in loadings on neurotraits, we label each group by dominant features (**fig. 6e**): Clinical features of neuropsychotype 1 reflected high pain qualities and negative affect, labeled as *vulnerable to pain;* neuropsychotype 2 indicated low pain qualities but high negative affect and low physical health, labeled *vulnerable to negative affect*; neuropsychotype 3 features indicated low pain quality and low negative affect despite higher levels of pain intensity, labeled *resilient*; neuropsychotype 4 features were near the average for all of the outcome measures, labeled *neutral*; neuropsychotype 5 indicated low levels of pain intensity and pain qualities, lower negative affect, and good physical health, labeled *protected*. These five neuropsychotypes when mapped on the network representation of the four chronic pain trait dimensions show patterns consistent with the labels associated with each type (**supplementary fig. 6**).

**Fig. 6.**
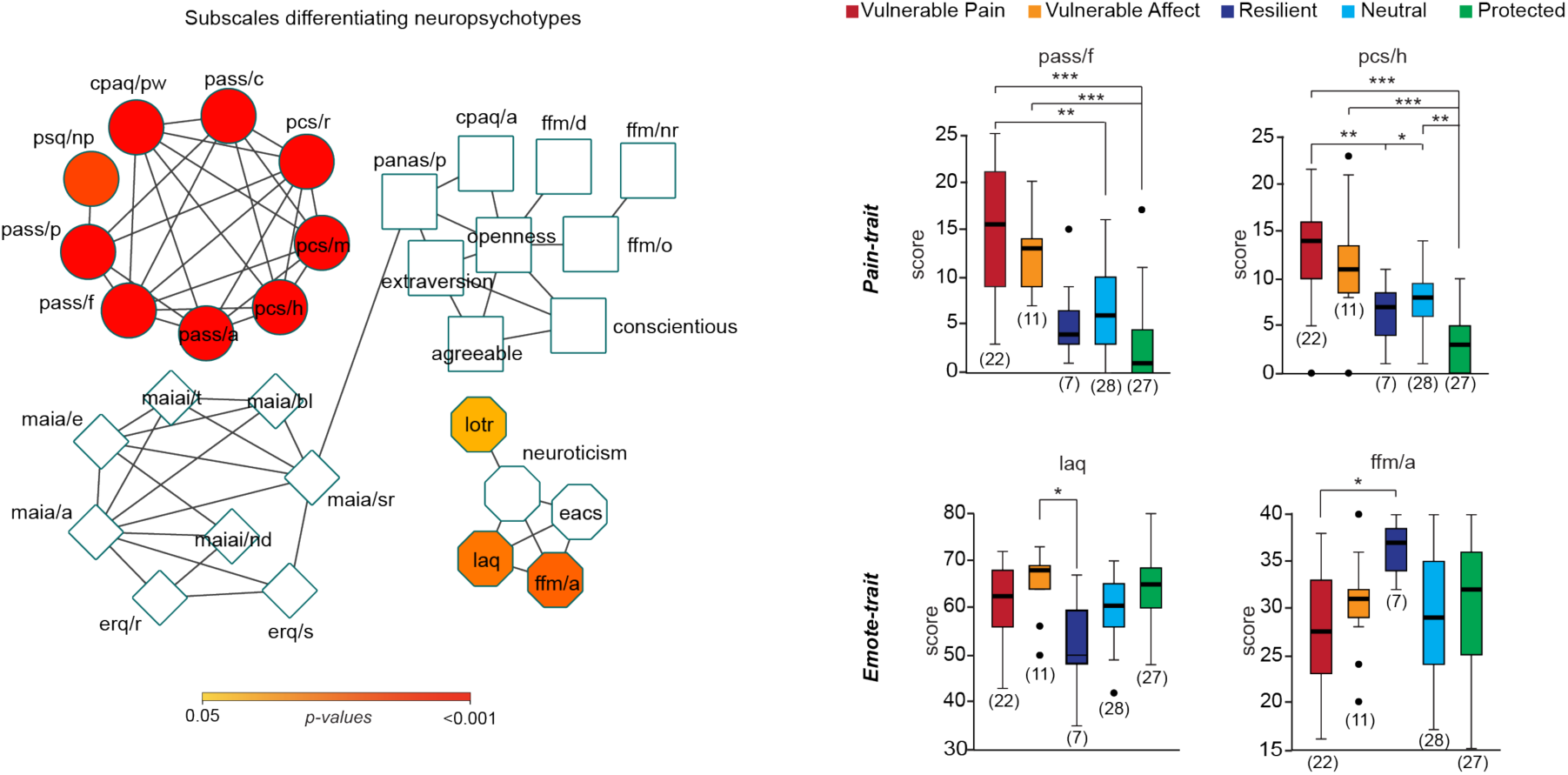
Relating traits and neuropsychotypes. **a.** Non-parametric comparisons indicated that neuropsychotypes differed on psychological factors loading on *Pain-trait* and *Emote-trait.* All subscales of *Pain-trait*, but not *Emote-trait*, survived *Bonferroni* correction for 36 comparisons (Group 1 and 2 combined, n=95). **b.** Post-hoc comparisons showed that the types vulnerable to pain and vulnerable to affect most strongly scored on subcales of the *Pain-trait* (e.g., pass/f and pcs/h) compared to the protected type. In contrast, the resilient type scored lower on the loss aversion (laq) questionnaire compared to the vulnerable to affect type and scored higher on the five facets of mindfulness acting with awareness (ffm/a) subscale compared to the vulnerable to pain type. Post-hoc comparisons were Bonferroni corrected for 10 comparisons * p < 0.05; ^**^ p < 0.01; ^***^ p < 0.001.

Here again, we tested the influence of pain duration on the neuropsychotypes and the results showed no differences in duration between the types (Kruskal-Wallis test; H=4.58, p=0.33).

We sought to validate these results using patients from Group 2, given that they expressed the same four chronic pain traits. Because fMRI data was not collected in these patients, we used the pain measurement to segregate these patients into the neuropsychotypes identified in Group 1 (**supplementary fig. 7a-e**). Concordant results were obtained in Group 2, as the five neuropsychotypes showed similar scoring on the 36 subscales (**supplementary fig. 7f**). The latter results also demonstrate that once neuropsychotypes are defined, then simple questionnaire outcomes are sufficient to pinpoint type memberships.

**Fig. 7.**
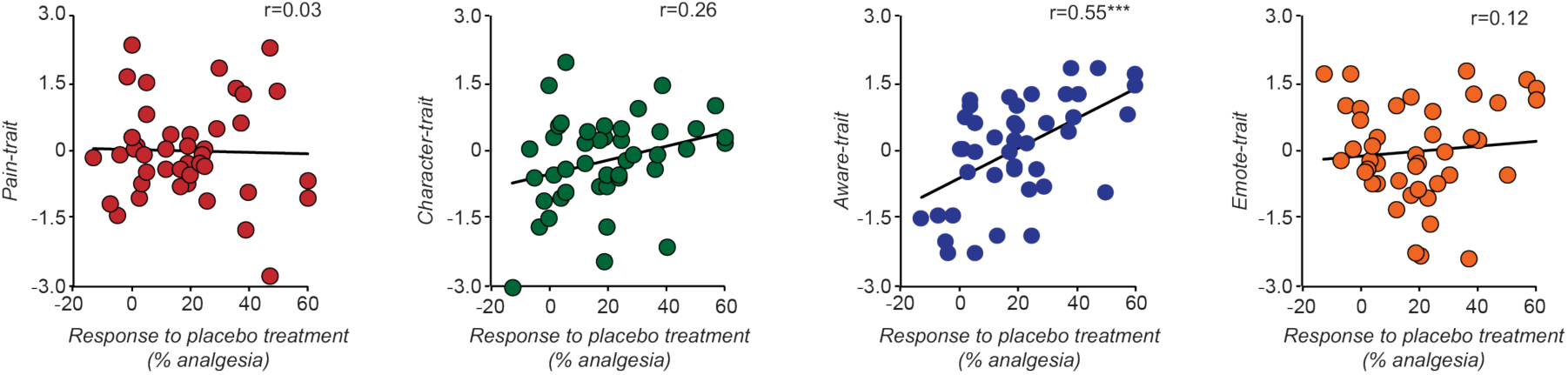
Traits predicting response to placebo pills. Only *Aware-trait 3* predicted the magnitude of the future placebo response (% analgesia). Correlation p-values were Bonferroni corrected for 4 comparisons *** p < 0.001.

Thus, **fig. 6a** shows the statistical comparisons between neuropsychotypes across all personality/psychology measures when considering all participants (Group 1 and Group 2 combined, n=95). The results show that *Pain-trait* conferred vulnerability and protection types, whereas *Emote-trait* determined resilience type (**fig. 6b**).

### Aware-trait 3 determined responses to placebo treatment

We further explored how the four chronic pain traits related to the response to placebo treatment during the RCT. We calculated the magnitude of placebo response as the strongest response (% change in pain) in either of the two treatment periods, as was previously done in (Vachon-Presseau et al., 2018). Interestingly, the 2 traits reflecting back pain properties of CBP, *Pain-trait* and *Emote-trait*, were unrelated to the placebo response. In contrast, *Aware-trait* was strongly predictive of the future magnitude of placebo response (**fig. 7**). The result is consistent with our earlier report (Vachon-Presseau et al., 2018) where we observed multiple subscales of MAIA being predictive of placebo response, and these all loaded onto *Aware-trait.* The current result highlights the fact that *chronic pain traits* unrelated to pain are still relevant when studying other characteristics of patients such as the likelihood of responding to a specific treatment.

### Socioeconomic properties of neuropsychotypes

Finally, we tested our hypothesis that socioeconomic status (SES) differentially impacts neuropsychotypes. We measured SES using self-reported income (US dollars earned yearly), race/ethnicity, gender, and years of education. First, we observed that income had the most robust impact on chronic pain traits, as patients reporting income greater than 25k expressed a lower level of *Pain-trait* (**fig. 8a**). This finding was replicated in Group 2 (**fig. 8b**). Moreover, the effect was not limited to psychological factors, as income was strongly linked with both positive and negative connections of *Neurotrait 1* and *Neurotrait 2* (**fig. 8c-d**), showing the influence of SES on the individual’s psychology and underlying neurophysiology. Income also showed differences between neuropsychotypes (borderline significant; Kruskal-Wallis test; H=9.16, p=0.057), with individuals who were vulnerable to pain type reporting less yearly earnings than their counterparts in the protected neuropsychotype. As displayed in **fig. 8f**, income mostly correlated with the subscales defining the traits of the neuropsychotypes (as displayed in **fig. 6a**), as lower income was linked with higher anxiety and catastrophizing, and higher income linked with higher optimism. Although years of education and race/ethnicity (but not gender) were strongly associated with income (**supplementary fig. 8**), the neuropsychotypes did not differ in gender, race/ethnicity, or years of education.

**Fig. 8.**
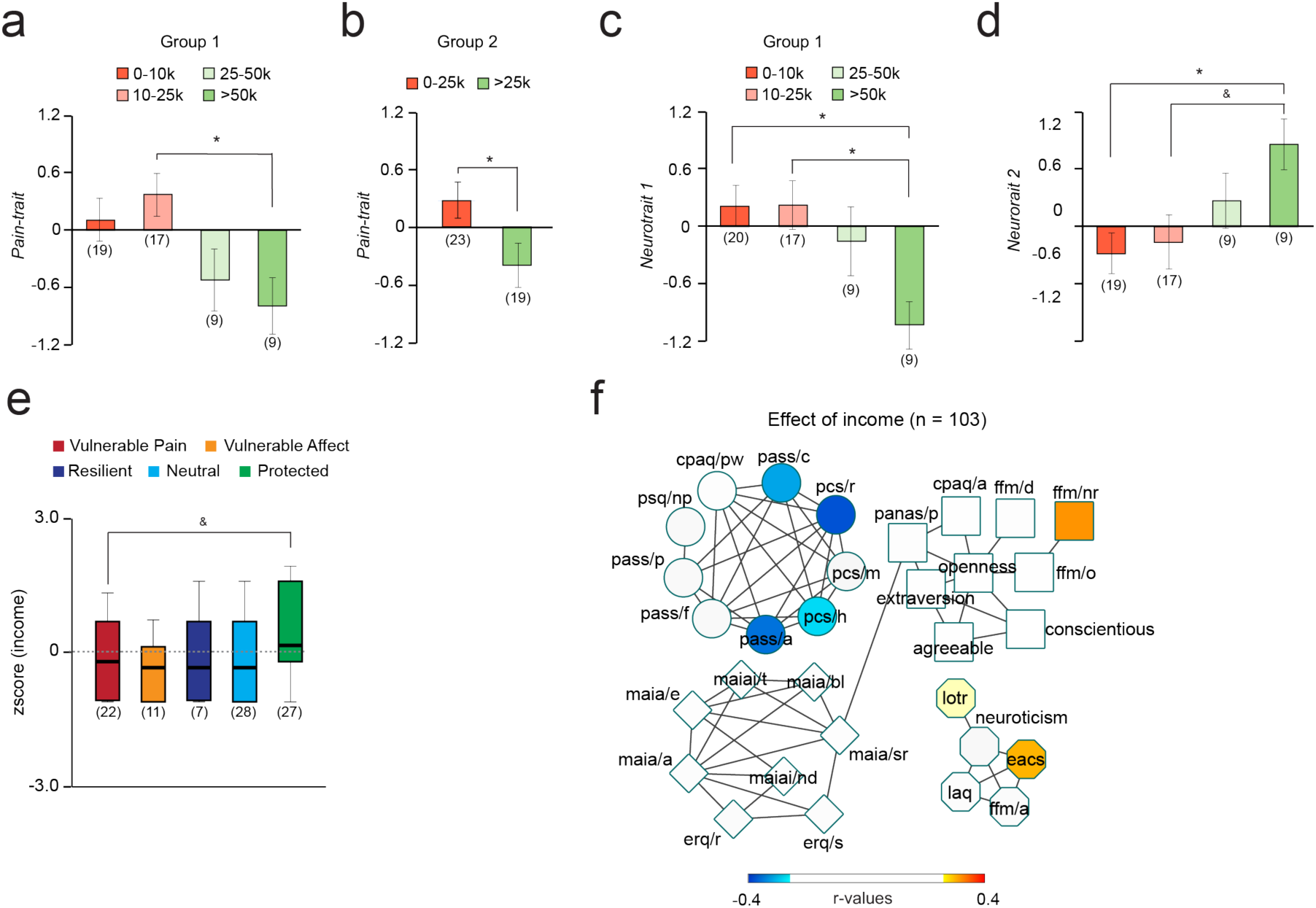
Influence of socioeconomic factors on traits, neurotraits and neuropsychotypes. **a.** Income differentiated patients on *Pain-trait* after controlling for ethnicity (while the inverse was not true), suggesting that socioeconomic status (rather than ethnicity) contributed to *Pain trait 1*. Annual income of >25k seemed to represent a socioeconomic threshold distinguishing vulnerability from protection. **b.** Applying the 25k threshold yielded similar results in Group 2. **c-d**. Self-reported income further differentiated neurotraits 1 and 2. **e.** Income differed between neuropsychotypes: vulnerable-to-pain types had lower income compared to the protected type (Group 1 and Group 2 combined). **f.** Spearman correlations were used to display subscales associated with income (only significant relations are displayed (uncorrected p<0.05). Post-hoc comparisons were Bonferroni corrected for 6 comparisons (**a,c,d**) and 10 comparisons (**e**) & *p* ≤ 0.1; * p < 0.05.

## Discussion

To our knowledge, this is the first comprehensive effort aimed towards defining the psychological/personality traits of chronic pain and integrating these properties with brain function to derive clinically-meaningful neuropsychotypes. We identified four chronic pain traits, which accounted for about 60% of the variance generated across a high-dimensional personality/psychology space (36 subscales from 13 questionnaires), validated in a separate sample. Only two traits modulated, in specific and opposite patterns, the pain characteristics. These were used to identify underlying brain circuitries (*neurotraits*), which were relatively stable over multiple weeks. Clustering the neurotraits and mapping them back into the traits-space and pain characteristics identified five neuropsychotypes, which were minimally dependent on pain intensity and independent of duration of back pain. Identified traits, neurotraits, and neuropsychotypes all showed dependence on socioeconomic factors – specifically, income – with the patients labeled as pain protected type showing higher SES status. Thus, our study provides a unified characterization of the chronic pain patient along psychological, neurological, and social dimensions, all underpinning specific traits and types.

Here, we advance a novel approach for deriving multivariate subtypes of chronic pain. By interrogating patients within a high-dimensional personality/psychology space – spanning the domains of attitudes towards pain (acceptance, catastrophizing, anxiety, sensitivity), interoceptive awareness, emotional regulation and attentional control, mindfulness skills, personality (extraversion, agreeableness, conscientiousness, neuroticism, and openness), optimism, motivation (loss aversion), and positive affective state – and then applying dimensionality reduction techniques (cluster and network analyses), we discover primary chronic pain traits and expound on their properties. Given that the dimensionality reduction methods used here depend on the input parameter space and the size of the population used, obtained results remain somewhat arbitrary, and one worries about missing critical dimensions. Nevertheless, our ability to a) validate the effects of traits on chronic pain measurements in an independent group of patients, b) show that the uncovered neurotraits were relatively stable across four consecutive brain scans, and c) demonstrate that changes in neurotraits mediated the changes in psychological factors and negative affect, altogether enhances confidence that the studied dimensions are robust, meaningful, and biologically-grounded.

### Traits of chronic pain

The current study is the first to systematically delineate the state space of the biopsychosocial (BPS) model for chronic pain. In this regard, the specific terminology that we invoke aims to highlight the limits of the BPS model as explored here. Although the BPS model has been advocated for over 30 years and is widely used in clinical pain settings (mainly in multidisciplinary pain management programs) (Gatchel et al., 2014; Haythornthwaite and Benrud-Larson, 2000; Jensen and Turk, 2014), it still remains a poorly-defined concept (Edwards et al., 2016): “The [BPS] model is rather vague about the specific pathways by which its elements interact and there are often no clear boundaries between categories of processes and constructs. Moreover, many of the explorations invoked by the biopsychosocial model to account for interindividual variability in pain-related outcomes are so multifaceted they are unfalsifiable by empirical research.”

Here, we begin delineating the BPS elements. Components of *Pain-trait* are all considered fundamental constituents of the BPS model. They have been repeatedly shown to be important contributors to chronic pain, and even seem to determine the analgesic efficacy of diverse therapies (topical or oral analgesics, cortisone, surgical outcomes, and psychosocial treatments like CBT) (Edwards et al., 2016). The next two factors, *Character-trait and Aware-trait*, reflect positive character, as well as awareness and emotional regulation traits, and showed no consistent relationship with back pain properties, although *Aware-trait* predicted response to placebo treatment. The fourth, *Emote-trait* factor, reflecting higher optimism, mindfulness capacities, and lower neuroticism and loss aversion was inversely related with negative affect. Thus, factors 2-4 seem to counterbalance the aversive toll of chronic pain and as such may be an important compensatory mechanism that patients use to tolerate the persistent CBP state (diminishing their perceived disability while living in chronic pain). From this viewpoint, *Emote-trait* specifically seems to offset and counteract the negative emotional impact of chronic pain, contributing to or perhaps even determining the resilient neuropsychotype.

Still, given that the approach is a first effort, it raises new and critical questions that require future study. First, there is a general consensus that the human population at large is composed of five broad personality traits, also known as the Five-Factor-Model (Goldberg, 1992). Our results show that most of these traits are clustered together in CBP (*Character-trait*), while other aspects (*Emote-trait*) become differentially related to chronic pain. Importantly, while in the population at large subjective well-being is observed in individuals high in extraversion and low in neuroticism (Ozer & Benet-Martnez 2006), in CBP these characteristics load on orthogonal factors (that is, they vary independently) and relate to distinct properties. How the CBP traits (and their interrelationships) change and congeal in the transition from acute to chronic pain is unknown and will be exciting to explore in the future. It will also be important to study the role of traits and individual differences in traits, in the context of treatments for chronic pain. Such knowledge would be important to know from a risk-management viewpoint. Concurrently, knowledge regarding malleability of these traits could provide targets for psychological or cognitive behavioral interventions. Second, we do not know the extent of invariance in the identified traits (and types) across different clinical chronic pain conditions (e.g. osteoarthritis, fibromyalgia, phantom pain, complex regional pain syndrome, pelvic pain, etc.), many of which are known to differ in their underlying biology (e.g., somatic vs. visceral conditions), or how the covariance (network architecture) between these traits may change after different kinds of treatments or therapies. Further studies exploring these issues are certainly warranted.

### Neurotraits of chronic pain

Data-driven methods provide an unbiased and optimal way to map distributed brain representations for complex phenomena (Kragel et al., 2018), and such an approach is especially appropriate when studying chronic pain since recent evidence indicates that pain patients exhibit a disruption of functional connectivity *throughout* the cortex (Mansour et al., 2016). We show that distributed patterns of functional connectivity successfully predicted *Pain-trait* and *Emote-trait* in left-out-patients. These relationships persisted over multiple weeks and, in part, mediated pain qualities and changes in negative affect of CBP. Not surprisingly, identified networks (*neurotraits*) involved brain regions not necessarily related to acute or chronic pain. The networks did still include a large number of connections involving the prefrontal cortex and the DMN, especially *Neurotype 1*, consistent with earlier evidence showing involvement of these regions in chronic pain and in catastrophizing (Kucyi et al., 2014) (Baliki et al., 2014; Loggia et al., 2013; Napadow et al., 2010).

Our results support the existence of multiple brain systems contributing and modulating chronic pain. These systems involved different circuitry representing a different mixture of psychological factors and personality traits. This is in accordance with the literature indicating that several systems may contribute to chronic pain. For instance, several studies have indicated that the DMN linked with pain catastrophizing determines pain intensity (Baliki et al., 2008; Baliki et al., 2014; Loggia et al., 2013), whereas the mesolimbic system associated with reward/punishment modulates pain and determines chronification (Baliki et al., 2012; Navratilova and Porreca, 2014; Ren et al., 2016; Schwartz et al., 2014; Woo et al., 2015). Accordingly, we demonstrated that psychological components of *Pain-trait* (pain catastrophizing, anxiety, acceptance) determining vulnerabitility or protection and *Emote-trait* (neuroticism, sensitivity to loss, optimism, and mindfulness capacities) determined resilience. These findings suggest multiple dimensions of personality interacting to determine chronic pain that have their own biology. Future studies should also incorporate neuroanatomical and structural brain properties, as these factors may further enhance the explanatory power of the neurotraits in relation to corresponding psychological/personality traits of CBP.

Providing a biologically-grounded model for chronic pain traits, which was missing so far, is itself relevant for several reasons. First, psychological contributors to pain are often considered intangible mental constructs that are rarely assessed in physical therapy clinics (Linton and Shaw, 2011). The demonstration that psychological contributors can be decoded from observable and quantifiable brain networks provides evidence that they are not uncontrollable noise but are instead biological determinants of the pathology. Secondly, the ResearchDomain Criterion (RDoC) framework suggests that diseases and mental illnesses should no longer be categorized from observable symptoms, but rather from the biology underlying the pathology (Insel et al., 2010; Insel, 2014). The RDoc suggests that biomarkers should be used for a better characterization of patients and a better understanding of the pathology, ultimately increasing the precision of diagnostics in the field of psychiatry. Similarly, different mechanisms promoting chronic pain can generate similar phenotypes (Denk et al., 2014) and the examination of biological mechanisms causing pain is necessary for identifying the right target to treat. Finally, better understanding of the biology of chronic pain in humans may improve translational research with animal models of chronic pain (Price et al., 2018; Vachon-Presseau et al., 2016).

### Neuropsychotypes of chronic pain

The field of personalized medicine is growing rapidly due to increased efforts in the development of objective measures derived from neuroimaging data that are capable of objectively determining *who* the individual experiencing a pathology is, and *how much* of that pathology they are dealing with (Duff et al., 2015; Woo et al., 2017). One recent study demonstrated that rsfcMRI can be used in major depression to generate homogeneous subtypes of patients sharing specific symptoms and commonalities in their responses to treatment (Drysdale et al., 2017). Using a similar strategy, we show that the expression of certain functional connectivity patterns derived from psychological determinants of chronic pain was capable of segregating CBP patients into homogeneous neuropsychotypes with similar clinical characteristics, providing clinically meaningful information about these patients’ personalities, coping skills, and overall well-being.

Given that the neuropsychotypes were derived only from the three neurotraits related to CBP properties, all five neuropsychotypes uniquely reflect various CBP properties and, thus, subdivide the trait-space in specific, chronic pain-related qualities. In future studies, we might expect to observe different distribution frequencies for neuropsychotypes across distinct chronic pain conditions while still preserving the specificity of responsiveness to therapies afforded by each neuropsychotype despite diverse symptoms and physiology. It is thus essential to validate the identified neuropsychotypes in other chronic pain conditions and also test the notion that distinct neuropsychotypes may be responsive to specific therapy regiments. After all, the concept of the BPS model is anchored on this notion and, thus, the neuropsychotypes should be viewed as the candidate types that define the BPS model and potentially link to treatment specificity.

A recent study, using a novel clustering approach and based on responses in more than 1.5 million participants to questionnaires measuring the personality traits of the Five-Factor model, finds robust evidence for four personality types in the population at large (Gerlach, 2018). The four types revealed by Gerlach et al. are denoted as: “average” type showing average scores on all traits; “Role Model” or “Resilient” type displaying socially desirable traits; and finally, the “Undercontrolled” and “Overcontrolled” types. Given that the starting trait-space studied by Gerlach et al. does not match the trait-space we identify for chronic pain, the resultant types in the two groups cannot show direct correspondences. However, there is a hint here that the chronic pain neuropsychotypes may map onto the population personality types or share substantial mutual information, such as resilience or protection. Thus, the chronic pain traits observed across our neuropsychotypes may *a priori* be present in the general population. Thus, the core types of resilience, overcontrolled-protected, and undercontrolled-vulnerable may already define *if* a healthy individual will develop chronic pain and *how* the chronic pain will be experienced. This too is an exciting idea that will require future studies.

### Socioeconomic influences on traits and types of chronic pain

Because chronic pain traits, neurotraits, and neuropsychotypes were all influenced by SES, one has to conclude that all are at least partially determined by environmental factors beyond the income, ethnicity, and education variables studied here to also include upbringing, culture, and politics. The differential influence of SES on neuropsychotypes, where the protected neuropsychotype had higher SES than the vulnerable neuropsychotype, enhances confidence in the consistency and interpretability of the identified types and highlights the potential utility of the neuropsychotypes in the realm of personalized medicine for chronic pain. Moreover, our results replicate previous research findings showing that lower incomes related with higher pain scores but also increased disability resulting from chronic pain (Rios and Zautra, 2011). Importantly, SES was most strongly linked with subscales defining the types of patients, as income correlated with components of *Pain-trait* (catastrophizing and anxiety of pain) and *Emote-pain 4* (optimism). Thus, the link between SES and our neuropsychotypes provides insights into the social component of the BPS model and further substantiates the idea that chronic pain experience is not only rooted in biology, but also intimately embedded in society.

## Conclusions

We advance a methodology for identifying traits and types in a specific chronic pain condition, CBP. The dimensions used to develop the trait space are globally relevant to all chronic pain conditions. By reserving the condition-specific pain characteristics, we could then use them to identify trait dimensions modulated by these parameters and identify related brain circuitry. We name these *neurotraits*. The identified *neurotraits* were then used to reveal neurophysiology-based CBP types, *neuropsychotypes*, that were mapped back into the trait- and pain characteristic-spaces to explore their properties. Besides providing a comprehensive specification of the BPS properties of CBP and showing the divergence of traits and types in CBP from the healthy population, the approach is applicable to all clinical chronic pain conditions, and thus, for the first time, provides tools and metrics that enable direct comparisons between such conditions regarding the neuropsychology of chronic pain. Moreover, the method can also be extended in the study of clinical negative mood conditions (anxiety, depression) and thus also enable comparisons between and across chronic pain and negative mood conditions. Additionally, we expect that the identified traits and neuropsychotypes will have clinical utility in personalized diagnostics and treatments.

## Methods

### Study design

This study was conducted in the setting of a clinical randomized controlled trial (RCT) specifically designed for assessing the placebo response (ClinicalTrials.gov: NCT02013427). The study consisted of 6 visits spread over approximately 8 weeks, including a baseline monitoring/screening period and two treatment periods, each followed by a washout period. The overall protocol included four scanning sessions collected before and after each treatment period. The design was set up to track placebo response in time and to test the likelihood of response to multiple administrations of placebo treatment. A full description of the study procedure on each visit, as well as the results of the placebo manipulation, can be found in (Vachon-Presseau et al., 2018). Here, we restricted our initial analyses to questionnaire and brain imaging data collected prior to placebo treatment. However, the brain imaging data acquired after the placebo treatment was used to test the stability of cross-validated patterns of functional connectivity and the effects of longitudinal changes on behavioral outcomes.

### Study Population

129 participants with chronic low back pain (CBP) were initially recruited from the general population via advertising in the community and clinical referrals via hospital databases. Patients were assessed for general eligibility via self-report using a screening intake form that the lab has used for years in other studies; this screening interview covered co-morbid health and psychological conditions, MRI safety, concomitant medication dosages and indications, pain levels/location/duration, current and previous illicit drug/alcohol use, litigation status, and overall willingness to be in a research study. To meet inclusion criteria, individuals had to be 18 years or older with a history of lower back pain for at least 6 months. This pain should have been neuropathic (radiculopathy confirmed by physical examination was required), with no evidence of additional co-morbid chronic pain, neurological, or psychiatric conditions. The enrolled patients had to report a pain level of at least 5/10 during the screening interview, and their average pain level from the smartphone app needed to be higher than 4/10 during the baseline rating period (explained below) before they were randomized into a treatment group. Finally, for safety precautions, clinical measurements taken at Visit 1 were required to be within the pre-specified healthy range (as determined by the standards utilized by Northwestern University Feinberg School of Medicine Laboratory Services Department) and all participants passed the MRI safety screening requirements at each scanning visit.

From the initial 129 chronic back pain (CBP) patients recruited in the study, 4 individuals were assessed for eligibility, but met exclusion criteria before consenting. Of the enrolled 125 patients, 63 chronic pain patients completed the study. One of these patients was removed from the brain imaging analyses because it failed quality control (see below). 47 additional patients completed all questionnaires from Visit 1, but did not complete the brain imaging session because they failed to meet the inclusion criteria at Visit 1 (BDI score > 19; n = 4), showed average pain levels lower than 4/10 during the 2-week baseline period between Visits 1 and 2 (n = 16), were claustrophobic or MRI incompatible (n = 9), had out-of-range vital signs or blood lab results (n = 6), or discontinued the study (e.g., were lost to follow-up, had increased pain, or no longer wanted to be in the study) (n=11). Participants were compensated $50 for each visit completed, and they were reimbursed up to $20 for travel and parking expenses if applicable.

### Ecological momentary assessments using phone app

Each patient’s pain was monitored electronically using an app designed specifically for the study. This app was used to track patients’ pain over time and to query them on their medication usage and mood; it could be accessed using either a smartphone or a website link on a computer. The app had a VAS scale with sliding bars: it asked participants to rate their current pain level from 0 (no pain) to 10 (worst imaginable). The app also included fields to indicate the participant’s assigned ID number, query if participants had taken any rescue medication at that time, and if they had taken the study medication. Participants were instructed to use the app twice a day, once in the morning and once at night. As an incentive, participants were compensated $0.25 for each rating they submitted, up to $0.50/day. This additional payment was given to them on the last visit of the trial to encourage compliance throughout the study. Submitted ratings were immediately sent to a secure server and both date- and time-stamped. Rating compliance was assessed by a separate program, which monitored whether the list of currently enrolled patients had provided the necessary ratings during the previous day. In the case that a patient omitted a rating, staff were alerted via an email. If patients missed more than 2 consecutive ratings (∼24 hours-worth), a member of the study team contacted them to remind them to use the app.

App rating data from all participants was pre-processed as follows. Although participants were asked to rate twice a day (and only compensated for this amount), many participants exceeded this number of app ratings in 24 hours due to over-compliance, reassessment of their pain, and/or cellular service problems. If pain ratings were entered within 30 minutes of each other, only the last rating was kept and taken as indicative of the participant’s final assessment of their pain levels at that time. Any additional ratings outside of this 30 minutes window were not considered duplicates and were kept as valid entries. Beside this cleaning process, no other changes were made to the ratings. In the instances where participants missed ratings, no attempts were made to interpolate or re-sample the data so that the temporal aspects of the ratings were left intact. The overall compliance of the phone ratings was on average 76%.

### Other chronic pain measurements

We assessed chronic pain using multiple measurements. First, we assessed the pain intensity using the phone app (see above), a numerical rating scale collected in the lab (NRS), and a verbal recall of the averaged pain experienced over the last week. Second, we assessed the qualities of pain using the McGill pain questionnaire (MPQ) affective and sensory subscales, and PainDetect. Third, we assessed mood with PANAS/negative subscale and the Beck Depression Inventory, Version 1a (BDI-1a). Finally, we assessed physical health using the 12-item short form survey (SF-12) physical subscale.

### Analysis of Questionnaire Data

A full list of all 13 questionnaires used to measure psychological and personality factors can be found in **supplementary table 2**. Data from these self-report measures was downloaded directly from REDCap as a CSV file and scored in Excel according to their references. Because all questionnaires were converted to an electronic format in order to be used in REDCap, an option to “skip” a question was provided if the participant did not feel comfortable answering a certain item. If more than 20% of the data from a given questionnaire (or questionnaire subscale, if applicable) was missing, the person’s data for the questionnaire was not scored; for all other missing data, the mean was used to fill in missing items (if the questionnaire had sub-scoring, the mean was calculated from the remaining items in the sub-dimension as opposed to the entire questionnaire); this approach is one of the most commonly used methods in data analysis (Pigott, 2001). It was utilized in order to conserve statistical power, given our relatively small sample size.

Data reduction was performed using two strategies. Firstly, a binarized undirected network was constructed using the strongest 10% correlation coefficients of the covariance matrix between the 38 subscales (i.e. network constructed at 0.1 density). Modules were identified using the Network analyser app in Cytoskape (http://www.cytoscape.org). Secondly, a principal component analysis was applied on the 38 subscales to extract four components explaining the above 63% of the variance. The components were orthogonalized using varimax rotation in SPSS version 21.

### Socioeconomic status

Self-reported income was measured with the following categories: 0-10K; >10<25k; >25<50k; >50K. Self-reported years of education was reported with a single number that ranged from 8-20. Race/ethnicity was also self-reported by the participant. There were a few missing data (patient skipped the question or preferred not to answer) for self-reported income, years of education, and ethnicity. In those cases, no attempt was made to fill those missing items. Because these missing values sometimes lead to analyses with different number of patients, the *n* are provided on each figure for each analysis.

### Brain imaging Protocol and Data Analysis

Brain imaging data was acquired with a Siemens Magnetom Prisma 3 Tesla. The entire procedure was completed in about 35 minutes, but an extra 25 minutes was allocated to install the patients in a comfortable position to keep their back pain at a minimum, and to re-acquire images if the data was contaminated by head motion.

High-resolution T1-weighted brain images were collected using integrated parallel imaging techniques (PAT; GRAPPA) representing receiver coil-based data acceleration methods. The acquisition parameters were: isometric voxel size = 1 × 1 × 1 mm, TR = 2300 ms, TE =2.40 ms, flip angle = 9°, acceleration factor of 2, base resolution 256, slices = 176, and field of view (FoV)= 256 mm. The encoding directions were from anterior to posterior, and the time of acquisition was 5 min 21sec.

Blood oxygen level-dependent (BOLD) contrast-sensitive T2*-weighted multiband accelerated echo-planar-images were acquired for resting-state fMRI scans. Multiband slice acceleration imaging acquires multiple slices simultaneously, which permits denser temporal sampling of fluctuations and improves the detection sensitivity to signal fluctuation. The acquisition parameters were: TR = 555 ms, TE = 22.00 ms, flip angle = 47°, base resolution = 104, 64 slices with a multiband acceleration factor of 8 (8 × 8 simultaneously-acquired slices) with interleaved ordering. High spatial resolution was obtained using isomorphic voxels of 2 × 2 × 2 mm, and signal-to-noise ratio was optimized by setting the field of view (FoV) to 208mm. Phase encoding direction was from posterior to anterior. The time of acquisition lasted 10 min 24 sec, during which 1110 volumes were collected. Patients were instructed to keep their eyes open and to remain as still as possible during acquisition. The procedure was repeated two times.

#### Preprocessing of functional images

The pre-processing was performed using FMRIB Software Library (FSL) and in-house software. The first 120 volumes of each functional dataset were removed in order to allow for magnetic field stabilization. The decision to remove this number of volumes was taken arbitrarily (it was not motivated upon examination of data) and we explored no other option. This left a total of 990 volumes for functional connectivity analyses. The effect of intermediate to large motion was initially removed using fsl_motion_outliers. Time series of BOLD signal were filtered with a Butterworth band-pass filter (0.008Hz<f<0.1Hz) and a non-linear spatial filter (using SUSAN tool from FSL; FWHM=5mm). Following this, we regressed the six parameters obtained by rigid body correction of head motion, global signal averaged over all voxels of the brain, white matter signal averaged over all voxels of eroded white matter region, and ventricular signal averaged over all voxels of eroded ventricle region. These nine vectors were filtered with the Butterworth band-pass filter before being regressed from the time series. Finally, noise reduction was completed with Multivariate Exploratory Linear Optimized Decomposition into Independent Components (MELODIC tool in FSL) that identified components in the time series that were most likely not representing neuronal activity. Components representing motion artifact were identified if the ratio between activated edge (one voxel) and all activated regions on a spatial component was >0.45, or if the ratio between activated white matter and ventricle and whole-brain white matter and ventricles was > 0.35. Moreover, noisy components were identified if the ratio between high frequency (0.05-0.1) and low frequency (0.008-0.05) was > 1. This ICA regression process was kept very conservative so that only components obviously related to motion or noise were removed.

The functional image registration was optimized according to a two-step procedure. All volumes of the functional images were averaged within each patient to generate a contrast image representative of the 990 volumes. This image was then linearly registered to the MNI template and averaged across patients to generate a common template specific to our CBP patients. Finally, all pre-processed functional images were non-linearly registered to this common template using FNIRT tool from FSL. The registered brains were visually inspected to ensure optimal registration.

On average, relative head motion was relatively low (mean frame displacement (FD) = 0.11; std 0.07 for the first rsfMRI run, and mean FD = 0.11; std 0.09 for the second run). Importantly, between-subject head motion was not related with any of the neurotraits (see below).

#### Parcellation scheme

The same parcellation scheme was used as in (Vachon-Presseau et al., 2018). The brain was divided into 264 spherical ROIs (5-mm radius) located at coordinates showing reliable activity across a set of tasks and from the center of gravity of cortical patches constructed from resting state functional connectivity (Power et al., 2011). In addition, 5-mm radius ROIs were manually added in the bilateral amygdala, anterior hippocampus, posterior hippocampus, and NAc. Linear Pearson correlations were performed on time courses extracted and averaged within each brain parcel. Given a collection of 272 parcels, time courses were extracted to calculate a 272×272 correlation matrix. These matrices were used for connectome-based predictive modeling, as described below.

#### Exclusion of participants

One patient from the no treatment arm was excluded from all rsfMRI analyses because of aberrant values in the correlation matrix (values were above 20 std from the mean). This subject was *a priori* rejected during initial quality check and was never included in any of the analyses (including the behavioral analyses. This subject was already identified as problematic and excluded in (Vachon-Presseau et al., 2018).

A second participant from the no treatment arm was excluded because of signal dropout in ROIs near the ventricle. This participant was included in our previous placebo study because we previously limited our analyses to 122 *apriority* selected ROIs (Vachon-Presseau et al., 2018). Here, we used the whole 272 ROIs and had to exclude that participant. The final sample size for brain imaging collected at visit 2 was therefore n = 61.

The longitudinal data included a total of n = 56 patients that passed quality check for each one of the four-brain imaging sessions.

### Connectome-based predictive modeling (CPM)

We used a validated data-driven protocol to predict behaviors (chronic pain trait scores) from brain connectivity measures between the ROIs of our parcellation scheme (i.e., the connectome). This approach was chosen because the CPM protocol has been argued to outperform most of the existing approaches (Shen et al., 2017). We first divided the patients into a training set (2/3 of the sample) and a testing set (1/3 of the sample). The training set was used to build the model, which was then applied to the testing set.

To train our model, we used robust regression between each one of the 36,856 connections with score on *Pain-trait (PC 1)* and *Emote-trait* (*PC 4)* (p < 0.05). We tested the stability of each using a leave-one-out procedure and selected only the edges correlating with the trait in each of the n-1 iterations. The end result was a number of stable edges positively or negatively correlating with *Pain-trait* score and *Emote-trait*. These edges’ strengths were summed to generate a single value, tracking each subject’s trait score.

To predict chronic pain traits in novel subjects, we extracted the summed strength of edges identified in the training set from the brain connectivity data in the testing set. This cross-validation procedure was repeated for the three folds, rotating and substituting the 1/3 of the patients used in the test set.

Two steps were used to ensure that motion did not influence the predictive patterns of brain connectivity. The first one was performed at the edge level, where removing the edges correlating with motion did not change the results. For instance, removing the edges related to motion at an arbitrary cutoff of p < 0.05 yielded very similar prediction accuracy in the test set (*neurotrait 1: r*^*2*^= 0.13, *p* = 0.004; *r*^*2*^ = 0.08, *p* = 0.03; *r*^*2*^ = 0.07*; p =* 0.04). Secondly, we tested the effect of motion at the individual level by regressing out inter-individual head motions. Thus, the pattern of functional connectivity (*neurotraits*) was uncorrelated with the mean head motion (Frame Displacement, FD) (all p values > 0.13) and regressing out the effect of the mean FD did not change the relationship between the *neurotrait 1* with *Pain-trait* (without covariates r = 0.83, p < 001; covarying for mean FD: r = 0.83, p < 001), *neurotrait 2 Pain-trait* (without covariates r = 0.91; p < 0.001; covarying for mean FD: r = 0.91, p < 001), and *Neurotrait 3* with *Emote-trait* (without covariates r = 0.87, p < 001; covarying for mean FD: r = 0.87, p < 001).

### Mediation analyses

All mediations were tested using the PROCESS macro implemented in SPSS. The macro allowed us to perform path analysis, standard mediation analyses, and sequential mediations. Here, significance was tested using 5000 permutations. The variable *neurotraits 1-2* used in the mediation analyses was generated using a linear regression entering *neurotraits 1* and *neurotraits 2* to explain *Pain-trait*. Both *neurotraits* explained independent variance and the variable *neurotraits 1-2* represent the predicted value from their joint contribution.

### Data availability

Data from our previous studies is already available on http://www.openpain.org/. The data used here is part of a longitudinal study that will generate more than one manuscript. The data will eventually be made available on openpain once these manuscripts are completed. Until then, data is available upon reasonable request.

## Acknowledgements

We are thankful to all Apkarian lab members who contributed to this study with their time and resources. We would also like to thank all the patients who participated in this study.

